# g.nome, A Transparent Bioinformatics Pipeline that Enables Differential Expression and Alternative Splicing Analysis by Non-Computational Biologists

**DOI:** 10.1101/2025.05.09.652286

**Authors:** Rut Bryl, Krystal C. Johnson, Xiongbin Kang, Tao Wang, Kit A. Fuhrman, Mark Kunitomi, Nora Kearns, David R. Corey

## Abstract

Reproducibility and accessibility are cardinal principles in the rapidly evolving field of bioinformatics. As the collection of biological data grows, proper use of pipelines to analyze datasets can become a bottleneck restricting efficient analysis. Biologists who collect data and test hypotheses may not have strong computational backgrounds and may not be able to fully understand the underlying strengths and weaknesses of computational approaches or fully exploit their data. Some data may be misunderstood and, perhaps more importantly, critical findings may remain unobserved. High throughput RNA sequencing (RNAseq) has advanced our understanding of transcriptomics across diverse applications. Here we introduce g.nome, a bioinformatics platform that integrates contemporary tools necessary for independent analysis. A user-friendly graphical interface simplifies running jobs and allows simplified analysis of different datasets by non-bioinformaticians. g.nome was used to analyze the consequences of localizing the critical RNAi factor argonaute (AGO) to nuclei of colorectal cancer cell line HCT116. Analysis using the pipeline facilitated the straightforward identification of splicing changes and the prioritization of these splicing changes for validation and further experimental analysis.

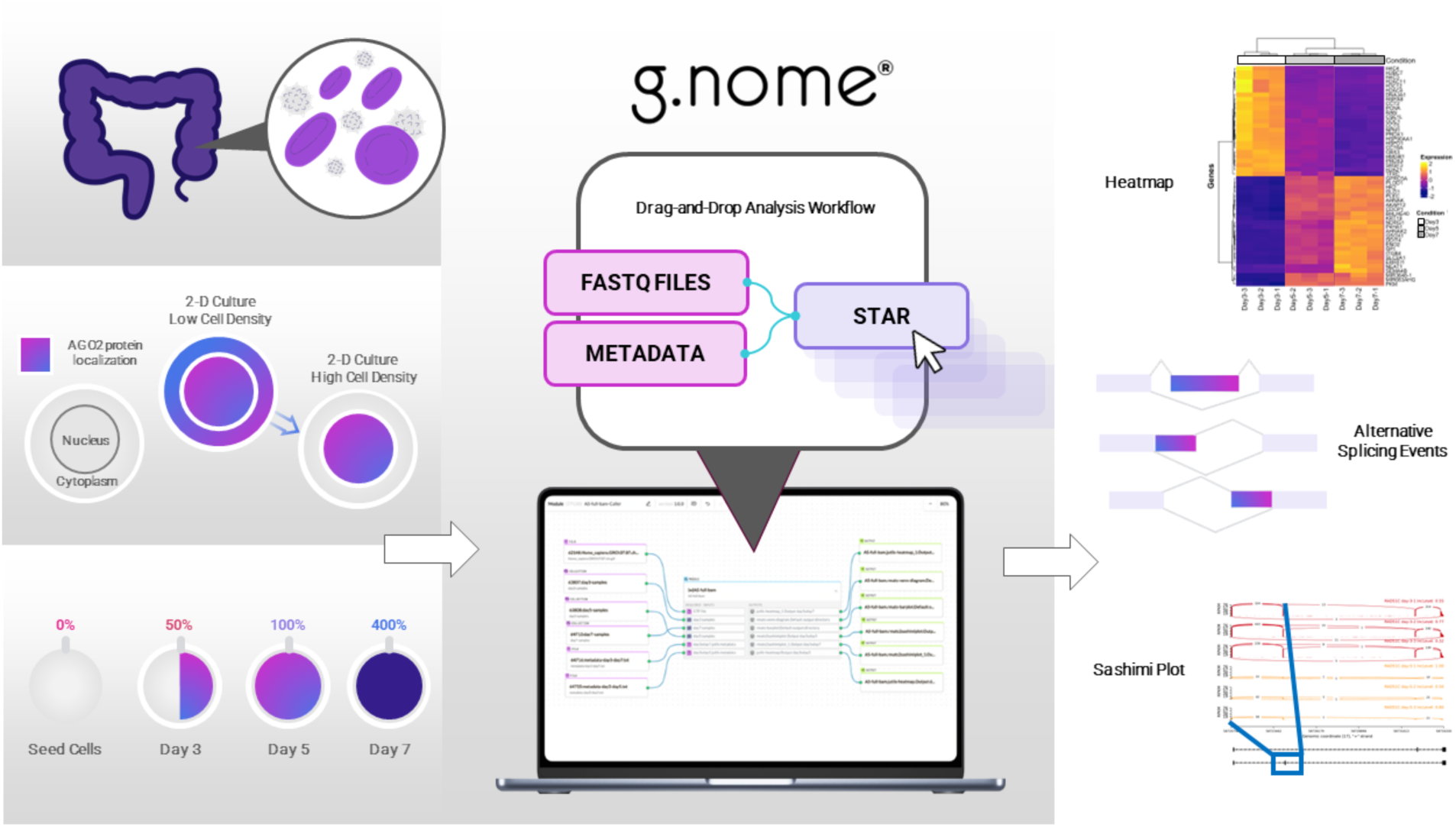

## Introduction

RNA sequencing (RNAseq) is becoming a routinely used tool for investigating gene expression and cellular regulation. RNAseq permits investigators to acquire large datasets and then compare how they differ as conditions change. These comparisons can identify the critical genes responsible for phenotypes and allow experimentalists to focus their efforts efficiently [1]. While the power of RNAseq to provide insights into biology is clear, translating that potential into useful insights is not straightforward. Most experimental biologists have a limited understanding of programming and computational analysis. They are, therefore, dependent on computational biologists to process RNAseq data and present it in a usable format for evaluation. In theory, this partnership can be an ideal combination of the strengths of different specialists [2].

In practice, over-burdened computational biologists may not have the time to fully engage and understand the biology questions related to the datasets. The experimental biologist may treat the computational biology as a black box whose details are not understood. Because these partnerships are challenging, information and insight can be lost in translation. The computational biologist may not adequately understand biology, leading to insufficient or improper analysis [3,4]. The experimental biologist may not understand how the data was acquired, leading to a misunderstanding of a dataset’s strengths and weaknesses.

To solve this misunderstanding, the g.nome application offers two input modalities that simplify bioinformatics for non-technical users. The first modality, g.nome Canvas, is a node-wire-language that containerizes bioinformatics tools as tiles that provides a clear way to track data flow through a workflow and can be run in series or parallel depeing on the workflow demands. Canvas’s tool tiles encapsulate the compute environment for each tool while exposing the various tool options to the user. The Canvas is currently engineered to run on Amazon Web Services but can be redirected to run on any scalable compute infrastructure. The second modality, Guided Workflows, is a form-based input method offering a series of simple questions and can be built on top of Canvas workflow or imported code.

Our overall goal was to develop a bioinformatics pipeline for analyzing data that can be used by experimental biologists who lack a programming background. To achieve this, we integrate an array of computational tools with the g.nome Canvas that are selected for specific strengths [5,6]. Starting with fastq RNAseq datasets, our pipeline systematically undergoes quality control, pre-processing, read alignment, post-alignment adjustments, transcript assembly, and abundance estimation to allow RNAseq alignments. These are incorporated into a transparent drag-and-drop interface that allows the user to intuitively understand which datasets are being analyzed, trace the flow of data through the pipeline and to easily change which comparisons are being made.

Our specific goal here was to demonstrate applying g.nome to the analysis of alternative splicing in a dynamic model system. Physiologically relevant models are necessary to bridge the gap between differential changes found in sequencing data and in vivo phenotypes. To enhance data interpretation and promote transparency in our analysis, we used the Jupyter Notebook as an extension of g.nome Canvas to generate a suite of visualizations tailored to our RNAseq results. Jupyter Notebook, an open-source platform, enables seamless integration of code execution, visualization, and narrative documentation, making it an ideal environment for exploratory data analysis [7].

We use colorectal cancer cells (HCT116) grown to high cell density as a model for solid colon tumors and applied g.nome to analyze their transcriptome [8]. When HCT116 cells are grown to high density, argonaute (AGO) proteins, which are normally evenly distributed between the cytoplasm and nucleus, shift localization to the nucleus. AGO proteins are the central effector proteins of RNA-induced silencing complexes (RISC) and are loaded with small noncoding RNAs known as microRNAs (miRNAs) to form miRISC [9-11]. miRISC recognizes and regulates the expression of target RNAs that contain sequence complementarity with the bound miRNA. Because the majority of the cellular AGO pool is enriched in the nucleus under these conditions, we hypothesize that this increase in nuclear concentration may decrease regulation of translation in the cytoplasm by miRNAs or promote AGO:miRNA-mediated recognition of intronic sequences and affect alternative splicing.

Previous work shows potential for miRISC-mediated regulation of alternative splicing [12-15] and testing the breadth of potential biological relevance demands optimal use of RNAseq datasets. Using g.nome, we have compared RNAseq data from cells grown to normal confluency (∼50-80% coverage of available surface area, AGO protein distributed between cytoplasm and nucleus) to cells grown to high confluency (∼400 %, or cells grown to form three or four layers, AGO protein primarily nuclear). The transparency and simple application of the alternative splicing and differential gene expression g.nome pipelines allowed rapid progress from analysis of data to hypothesis generation and finally to experimental tests [4,6].

## Materials and Methods

### Implementation

g.nome is a cloud-based platform offering end-to-end solutions for bioinformatics analysis, including data management, pipeline execution, interactive coding via Jupyter Notebooks, and result visualization. A key feature is its **Guided Workflows**—a graphical user interface (GUI) that enables no-code execution of Nextflow pipelines.

In this example, the **<u>nf-core/rnaseq v3.14.0</u>** pipeline was launched using the g.nome Guided Workflow interface. Maintained by a global community of bioinformaticians, nf-core/rnaseq is one of the most popular workflows in the nf-core repository, with over 1,000 GitHub stars. The pipeline is modular, allowing for easy tool-level customization, and each process is encapsulated within a public Docker container to ensure portability and reproducibility. Reference genomes can be automatically retrieved from ngi-igenomes, streamlining setup.

The nf-core/rnaseq pipeline is open-source and available to run independently off g.nome, provided a user has sufficient compute resources. However, it requires the computational expertise to configure and execute. The Guided Workflow feature in g.nome simplifies pipeline configuration for biologists by translating parameter settings into intuitive form-fill steps—eliminating the need for direct coding.

### RNAseq-based differential expression and alternative splicing analysis using g.nome

*Differential Expression Analysis*. Differential gene expression analysis was conducted using **DESeq2** (v1.34.0). The gene count matrix and sample metadata were used as inputs, with d3 as the control group. Differentially expressed genes were identified using a p-value cutoff of 0.05 and a fold-change cutoff of 0.75. Results were output to a specified directory.

*Gene Set Enrichment Analysis (GSEA)*. Gene set enrichment analysis was performed using **GSEA** with human-specific gene annotations. The DESeq2 dataset and associated metadata were used to identify enriched gene sets, applying a fold-change cutoff of 0.75 and a p-value cutoff of 0.05.

### Portability to AWS Cloud and Computing Cost

The pipeline nf-core / rnaseq is run on AWS batch where the compute environments are created with the following settings: on-demand instance, max CPUs 1000, NVMe enabled, and fusion mounts enabled. Instance types are chosen from the following list (EC2 instances with NVMe): m6id.large, r6id.24xlarge, c6id.12xlarge, m6id.24xlarge, r6id.large, r6id.2xlarge, c6id.32xlarge, m6id.8xlarge, r6id.32xlarge, c6id.8xlarge, r6id.xlarge, m6id.12xlarge, m6id.metal, c6id.24xlarge, r6id.metal, r6id.12xlarge, m6id.16xlarge, r6id.4xlarge, m6id.32xlarge, c6id.metal, c6id.16xlarge, c6id.4xlarge, r6id.8xlarge, m6id.2xlarge, m6id.xlarge, c6id.2xlarge, c6id.large, r6id.16xlarge, m6id.4xlarge, c6id.xlarge with the allocation strategy ‘best fit’. The computation is run in AWS region us-east-1. The pipeline runs are launched with Nextflow version to 24.04.4, build 5917 by adding export NXF_VER = 24.04.4 to the pre-run script.

### Scalability

Nextflow uses reactive programming to stream data dynamically between processes. As soon as data from one process becomes available it passes on to the next without waiting for the entire dataset to complete a process before starting the next one. Each process runs in an isolated, containerized environment which enables easier distribution across nodes in a cloud environment.

### Use of g.nome pipeline for alternative splicing analysis by non-bioinformaticians

Sequencing data and the reference genome alignment file were saved in a folder on the user’s computer. The raw sequencing files and human genome assembly were uploaded into the “Data” folder on g.nome. Under “My Projects”, the “Alternative Splicing and Gene Counts” ➔ “Go to Data” was selected **(Suppl. Fig. 1A, B)**. Then, in the upper right corner, “Add” ➔ “Collection” was selected to create a batch collection of relevant data. This collection was named, “Day3-Forward”, and the search bar was used to look up the relevant files with “Day3” and “R1” in the file names for forward reads for each of the three biological replicates from the Day 3 time point.

After these 3 files were selected, “Create” was clicked **(Suppl. Fig. 1C)**. This process was repeated to create batch collections for “Day3-Reverse” by selecting the three files that contained “Day3” and “R2” for reverse reads. Then this process was repeated to create batch collections for the other time points: “Day5-Forward”, “Day5-Reverse”, “Day7-Forward”, and “Day7-Reverse”. Under the Modules tab at the top center of the page, the alternative splicing pipeline module, “Runner AS and Genecount,” and “Edit Workflow in Builder” was selected **(Suppl. Fig. 1D)**.

Within the splicing module, there are empty cells for “requested inputs”. The orange circle on the left side of “Sample 1 – name” was dragged to the left into the open workspace area. The “String Value” was edited to be the sample name, “Day3”. This was repeated to name the other sample “Day7”. Then, on the right side of the screen, “Data” was selected, and the search bar was used to look up the batch collections created previously for sequencing data separated by time point and forward/reverse reads. I clicked and dragged and dropped the forward and reverse batch collections for “Day3” and “Day7” into the right-hand workspace. To designate the input for sample 1 on the splicing module, the orange circle on the “Sample 1 – fwd reads” required input was dragged to connect to the open white circle on “Day3-Forward” data collection. This was repeated for the reverse collection for Day3 and for forward and reverse collections for “Sample 2” inputs. Finally, on the right side panel, the “Run” button was selected **(Suppl. Fig. 1E)**.

The edited workflow can be saved, and the progress of the run can be monitored within the “Runs” tab at the top center panel **(Suppl. Fig. 1F)**. The down arrow on the run can be selected to reveal a dropdown menu with output files such as quality control measurements and summary graphs displaying analyzed data.

### Use of g.nome pipeline for differential gene expression analysis by non-bioinformaticians

To ease the setup and running of bioinformatic runs, g.nome allows for an additional layer of abstraction on top of Canvas Built workflows called Guided Workflows. In this case, the Guided Workflows executes the same underlying Canvas workflow but enables users to enter input values such as files, samples sheets or tool options in a simplified web form. Uploading the raw sequencing data and accessing the project was analogical to alternative splicing analysis (see “Usage of g.nome pipeline for alternative splicing analysis by non-bioinformaticians”).

Under “Workflows” tab, “Guided Workflows” button was clicked and the differential gene expression pipeline module “RNA-seq Preprocessing & Analysis” and “Run Guided Workflow” was selected (**Suppl. Fig. 2A**). The run was labeled and collections “Day357-Forward” and “Day357-Reverse” were selected as sample FASTQs (**Suppl. Fig. 2B, 2C**). The FASTQ files were then paired, according to their biological replicates (**Suppl. Fig. 2D**). Next, the samples were grouped – “Manual Edit” button was selected, and control group and group lists were defined. Finally, samples were matched with their respective group (**Suppl. Fig. 2E**). Reference genome was selected as “Workflow Supplied reference data” and “GRCh38” (**Suppl. Fig. 2F**). A read count method was “FeatureCounts” (**Suppl. Fig. 2G**). Optional parameters were left as default (**Suppl. Fig. 2H**). Before launching the run, the user is provided with the summary of the chosen parameters (**Suppl. Fig. 2I**).

The run can be monitored in the “Runs” tab (**Suppl. Fig. 2J**). The name of the run was then clicked, which takes the user to the outputs, where analyzed data can be accessed (**Suppl. Fig. 2K**). Basic parameters such as p-value, log fold-change, reference genome as well as sample metadata and FASTQ files used can be viewed in “Parameters” tab (**Suppl. Fig. 2L**).

### Preparation of cytoplasmic and nuclear extract from cell culture

Wild-type HCT116 cells were cultured in McCoy’s 5A medium supplemented with 10% FBS. Cells were incubated at 37°C in 5% CO 2 and passed when 70–80% confluent. Cells were seeded at 100,000 cells per well into 6-well plates on Day 0, and 8 ml of fresh media was supplied every day starting on Day 2 through Day 7. Cells were detached with trypsin, harvested, and counted with trypan blue staining (TC20 Automated Cell Counter; Bio-Rad). Subcellular fractionation to isolate cytoplasm and nuclear lysate extract was similar as previously described [14, 16, 17, 18]. Keeping the samples on ice, using ice-cold buffers, and using gentle pipetting is critical for maintaining the integrity of the nuclear membrane before and after cytoplasmic lysis.

### Preparation of whole transcriptome RNAseq libraries

Total RNA was extracted from whole cells grown in 2D culture for 3 days (50% confluency), 5 days (250% confluency), and 7 days (300–400% confluency) with TRIzol. The RNA quality was measured using the Agilent 2100 Bioanalyzer system with the RNA Nano chip kit (catalog# 5067-1511), and the RNA concentration was measured using the Nanodrop 2000 spectrophotometer by Thermo Scientific. Roche Kapa RNA HyperPrep Kits with RiboErase (HMR) (Catalog #KK8561) were used to generate the RNA libraries. The procedure followed the manufacturer’s instructions. The RNA input amount for each library preparation was 1.0 μg. The workflow began with rRNA depletion, followed by RNAse H and DNase treatment. Subsequently, fragmentation was carried out at 94 ◦C for 5 min. The cleaved RNA fragments were then reverse transcribed into first strand cDNA synthesis using reverse transcriptase and random primers. Second strand cDNA synthesis and A-tailing were performed, with incorporation of dUTP into the second cDNA strand. UMI adapters synthesized by IDT (Integrated DNA Technology) were used during the ligation process. The cDNA fragments underwent purification and enrichment via 9 to 10 cycles of PCR amplification to create the final cDNA library. Library quality was verified using the Agilent 2100 Bioanalyzer, and the concentration was measured using the picogreen method. The nine RNA-seq libraries were sequenced on Illumina NovaSeq 6000 sequencer platform using the PE-150 bp (paired-end) protocol.

## Results

### Experimental Design

We have previously observed that the localization of AGO protein can change as cell density increases [8]. In these experiments, we prepared RNA from HCT116 colorectal cancer-derived cells. When cells were grown to ∼50% confluence, AGO2 was distributed between cytoplasm and nucleus. When cells were grown to ∼250%, or to ∼400% confluence (equivalent to four layers of cells), AGO was predominantly nuclear. Cell confluence was determined by counting the number of cells in conjunction with the known surface area of the culture dish wells. The localization of AGO to cell nuclei suggestion that the gene regulatory pathways controlled by AGO protein would be perturbed. We used RNAseq to evaluate changes in gene expression.

RNA for RNAseq was extracted either three (∼50% confluence), five (250% confluence), or seven (∼400% confluence) days after seeding cells and used for whole transcriptome sequencing (**Fig. 1A**). Localization of AGO2 protein to the nuclei of cells used for sequencing was confirmed by western analysis of cytoplasmic and nuclear fractions (**Fig. 1B**). There are three other AGO proteins in human cells, AGO1, AGO3, and AGO4, these proteins also localized to cell nuclei [8]. The purity of the fractions was confirmed by detection of ß-tubulin as a marker of cytoplasm and histone H3 as a marker for nuclei. At 50% confluence most AGO2 was nuclear but a substantial fraction remained cytoplasmic. At 400 % confluence AGO2 was almost entirely nuclear. All RNA preparations and RNAseq analyses were performed in triplicate. The RNA libraries were sequenced on the Illumina Novaseq 6000 sequencer platform using the PE-150 (pair-end) protocol.

**Figure 1.**
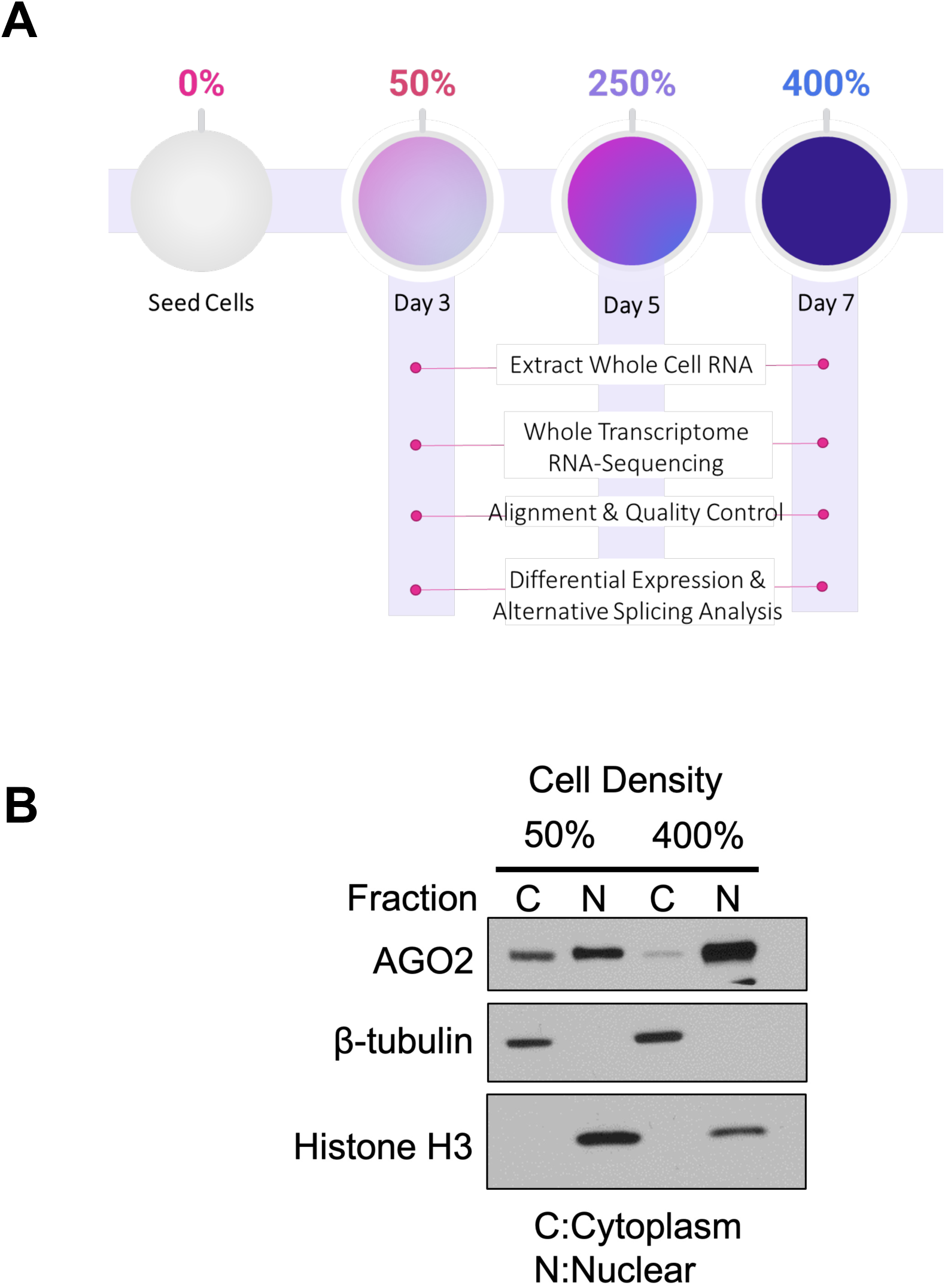
Using high cell density model system to drive AGO2 enrichment in the nucleus and explore consequences on alternative splicing. (A) Overview of RNA analysis workflow. RNA collection at different cell densities for whole transcriptome RNA-sequencing and g.nome module used for alignment, quality control, differential expression, and alternative splicing analysis. (B) Western analysis of cytoplasmic/nuclear distribution of AGO2 at 50% (Day 3) and 400% (Day 7) cell density. ß-tubulin and Histone H3 are markers for cytoplasmic and nuclear purity, respectively.

### Pipeline Overview and Data Processing

The goal of g.nome is to make RNA-seq analysis more straightforward for users, especially those with less bioinformatics or coding experience. It simplifies building pipelines because it removes the need for coding expertise and provides a visual framework for tracking data flow through various bioinformatic tools. Pipelines created using g.nome, including those described in this manuscript, are distributable, robust, and reproducible, and should operate on most computing environments. **Fig. 2A** presents a basic view of the g.nome alternative splicing module in a simplified interface intended for non-bioinformaticians. The module connects the inputs, where user can define their forward and reverse reads collections (FASTQ files), sample metadata, and reference genome, with the outputs - rMATs jobs.

**Figure 2.**
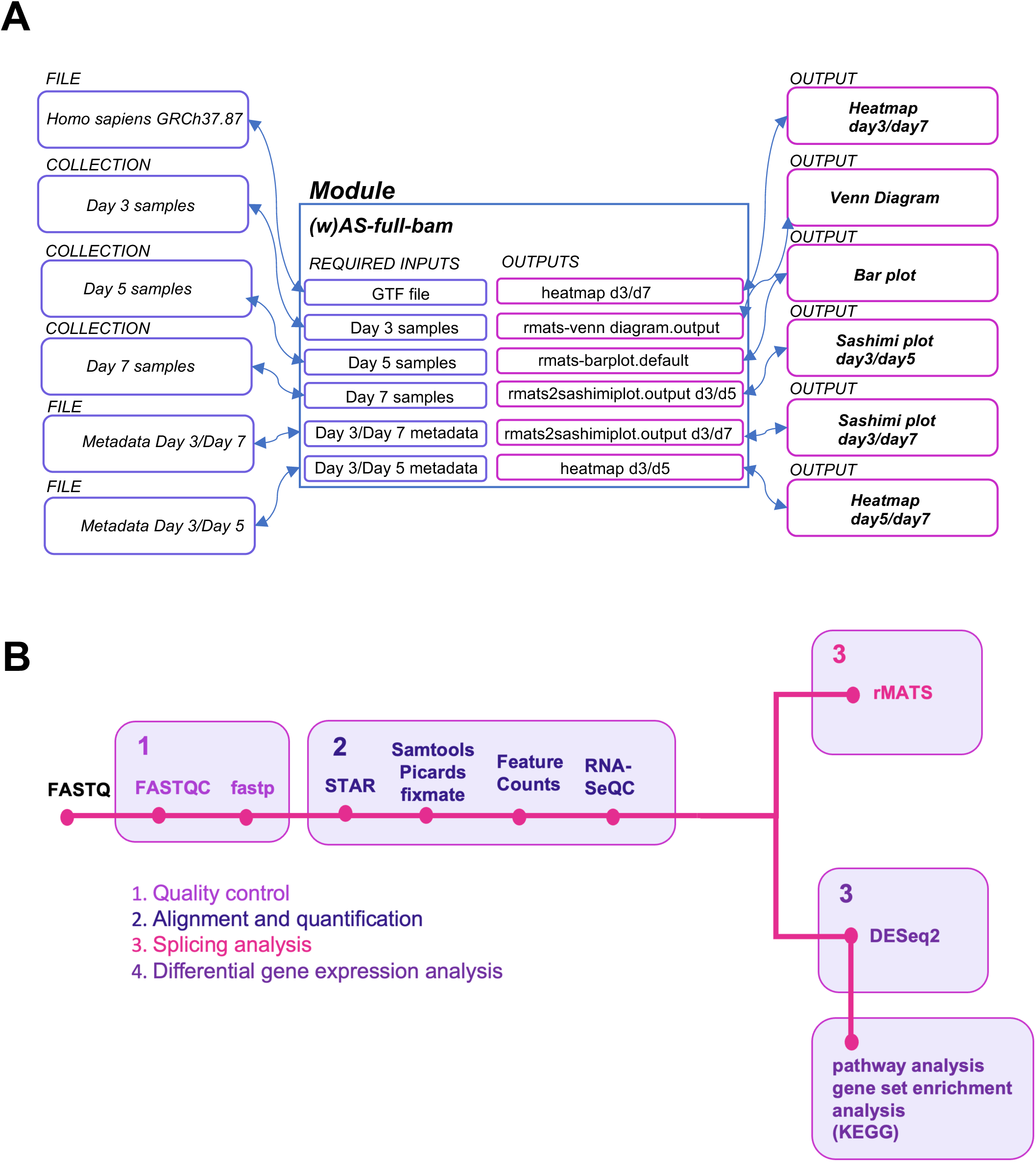
Overview of differential gene expression and alternative splicing analyses using g.nome pipeline. (A) Basic view of g.nome alternative splicing module for biologists to run analysis of Day 3, Day 5, and Day 7 samples and rMATS jobs in simplified interface. (B) Workflow of differential gene expression and alternative splicing analysis of RNA sequencing data using g.nome.

After data acquisition, the first step for g.nome is to select the pipeline and choose parameters. g.nome contains multiple different premade pipelines for different types of analyses. G.nome also includes graphical user interface that enables users to build custom pipelines that can then be used more generally by subsequent users. In this case, we built an RNAseq pipeline for analysis of differential gene expression and alternative splicing.

Following the acquisition of RNA-seq data from cells at different confluence, we selected a differential gene expression/alternative splicing pipeline with the ability to set optional parameters (**Fig. 2B**). After initiating analyses, quality control of reads is accomplished by FastQC (a tool that examines the raw data from fastq files and provides a summary of statistics that includes base quality, read length, G/C content, repetitive content, and adaptor content).

During library preparation, adaptor sequences are added to facilitate RNAseq. After obtaining a readout of quality with FastQC, the data is processed using fastp, which includes trimming of the adaptor sequences and quality filtering that removes reads that do not meet criteria, such as minimum length, or minimum quality score. This ensures that only high-quality data is used in downstream analyses. FastQC is then run again to determine whether quality has improved.

After quality control, alignment to the reference genome is performed using the STAR aligner, and the aligned reads are sorted with Samtools sort and Picards FixMateInformation. FeatureCounts is then used for transcript assembly and abundance estimation, while RNA-SeQC assesses the quality of alignments post-assembly. For alternative splicing analysis, rMATS identifies five types alternative splicing events, with rmats2sashimi providing visualization via Sashimi plots. The process of using the outputs to run Sashimiplot visualizations was streamlined to provide comprehensive results as opposed to “per splicing event” data. For differential gene expression analysis, DESeq2 was used, followed by pathway analysis from Kyoto Encyclopedia of Genes and Genomes (KEGG) (**Fig. 2B**).

### Global RNA Expression Analysis Across Cell Densities

We first used g.nome to examine which genes were differentially expressed between Day 3, 5 and 7. Principal component analysis (PCA) indicated that the triplicate datasets were closely matched (**Fig. 3A**). We then applied the MA (M:log ratio, A:mean average) plot to visualize expression level and differential expression of genes (**Fig. 3B**). Individual genes that combine changes that are both large (≥ log_2_ *FC* (1.5)) and significant (p value ≤ 0.05) were visualized by heatmap and volcano plot (**Fig. 3C, D**). Our analysis revealed significant changes in global RNA expression as cell density increased to 250 or 400 %. More genes were activated than were repressed, consistent with the conclusion that miRNAs represses gene expression and that repression is released when AGO moves from being a cytoplasmic protein to being a nuclear protein that is no longer available to fulfill its canonical role.

**Figure 3.**
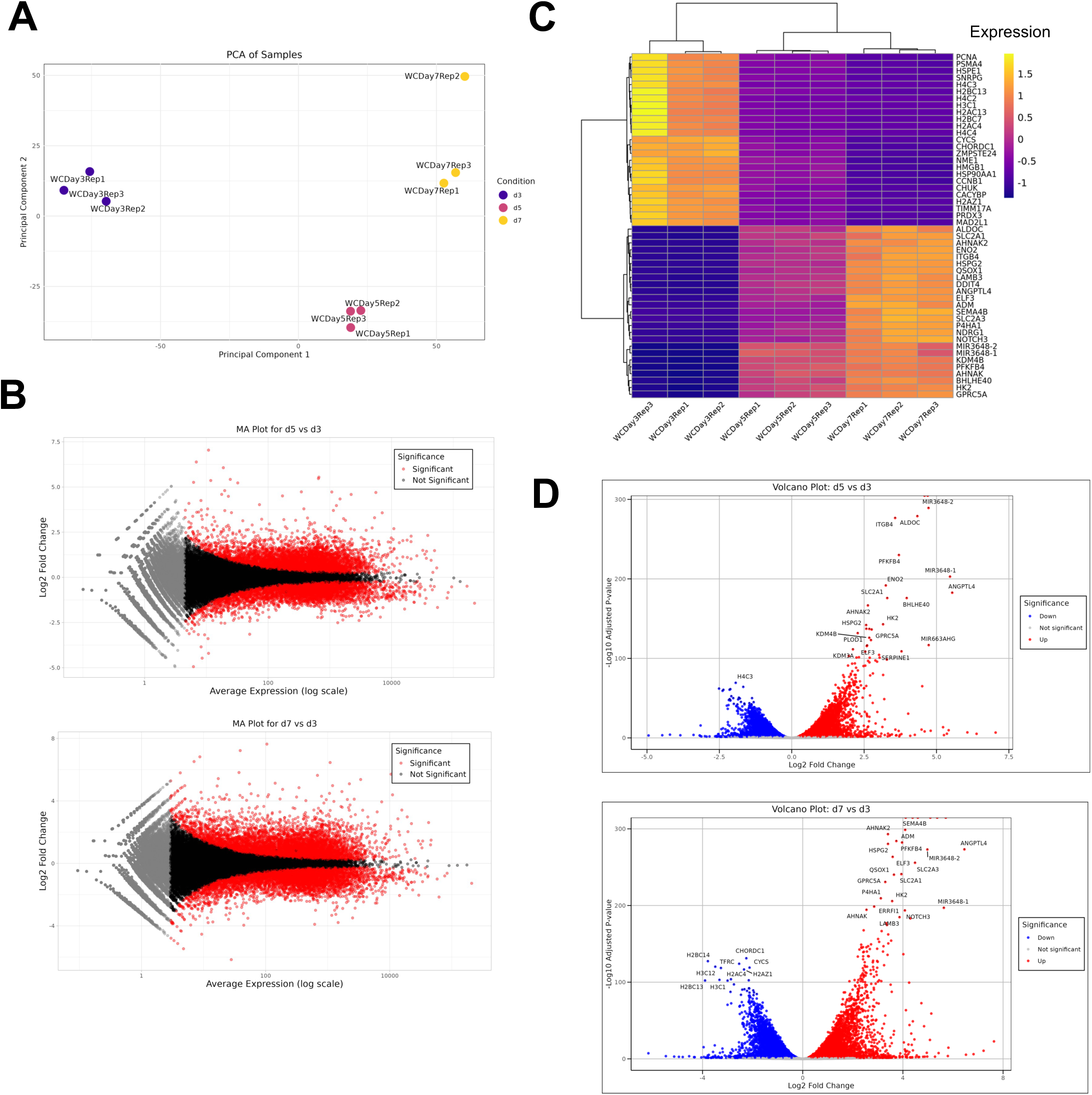
Global differential steady-state RNA expression analysis at different cell densities. (A) Principal Component Analysis of three replicates per cell density condition shown: Day 3 (50% cell density), Day 5 (250% cell density), and Day 7 (400% cell density). (B) MA-plot showing differential expression of each gene against its average expression across all samples. Significance shown for Day 5 (250% cell density) relative to Day 3 (50% cell density) and Day 7 (400% cell density) relative to Day 3 (50% cell density). (C) Heat map showing top 50 up- or downregulated genes across three cell density time points (Day 3, Day 5, Day 7) and their biological replicates (N=3). (D) Volcano plots showing genes significantly downregulated (blue) and upregulated (red) in Day 5 (250% cell density) relative to Day 3 (50% cell density) and Day 7 (400% cell density) relative to Day 3 (50% cell density).

### Pathway and Biological Process Enrichment

Many biologists benefit from, not only differential expression analysis, but also putting the data into the context of biological pathways. To provide these types of analyses, the pipeline developed on g.nome contains connections between the differential expression data outputs and modules to perform both pathway analysis and gene set enrichment analysis. These analyses provide visualizations to facilitate interpretation of data (**Fig. 4**). When HCT116 cells are grown to high density, we observed a shift in the localization of AGO2 to the nucleus [8]. One of our goals was to use the RNAseq data to investigate how cellular processes change when AGO2 is enriched in the nucleus. For our data, pathway analysis revealed a set of biological pathways that were significantly altered at high cell density. Notably, at 400% cell density, certain pathways were enriched, indicating a density-dependent regulation of cellular processes (**Fig. 4**). Other changing pathways, including ribosome biogenesis and mitochondrial translation, are consistent with increased cell crowding and associated lower growth rates.

**Figure 4.**
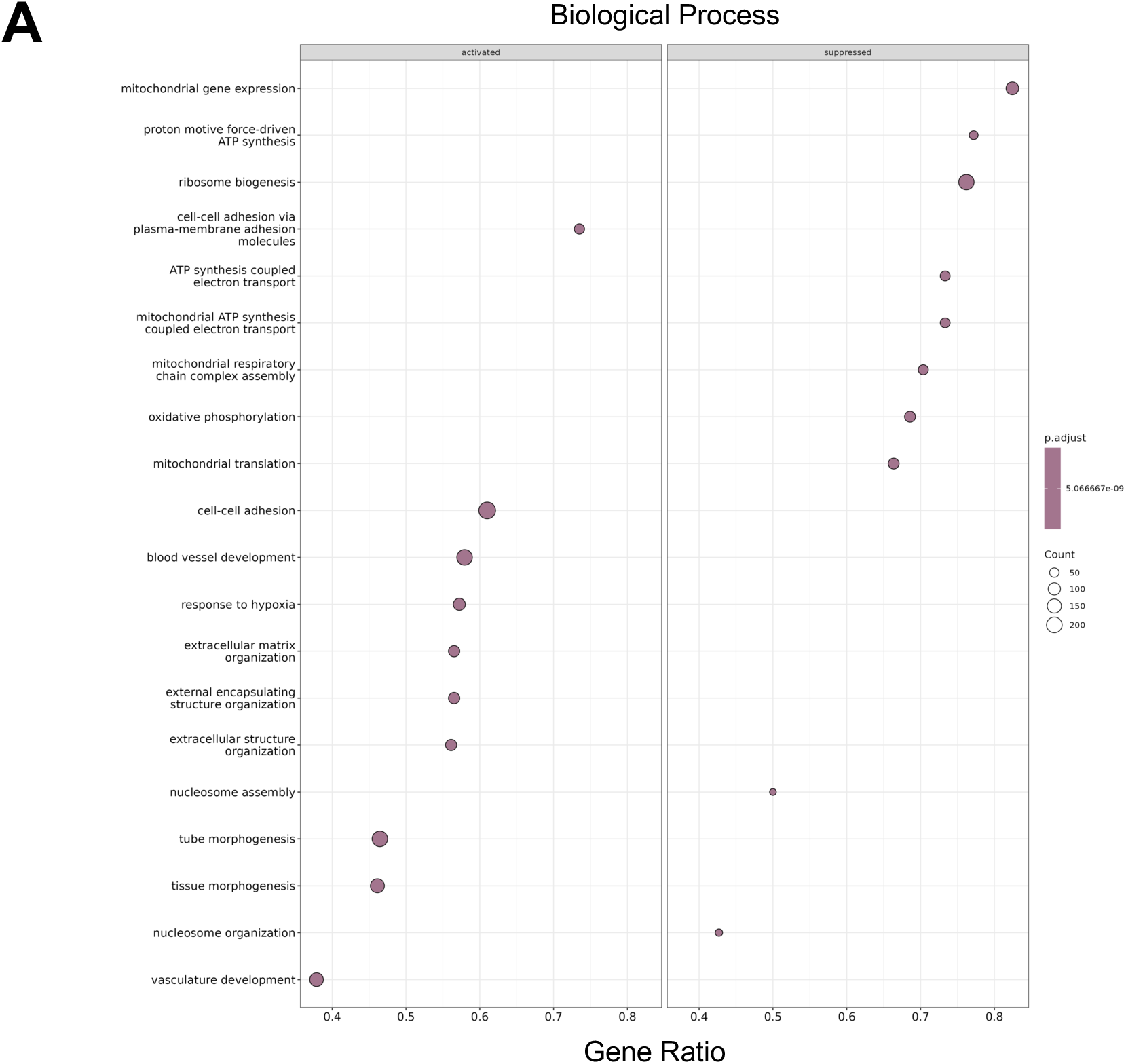
Differentially expressed pathways correlated with cell density and nuclear AGO2 enrichment. (A) Pathway analysis from the Kyoto Encyclopedia of Genes and Genomes (KEGG) highlighting pathways changing on Day 7 (400% cell density) relative to Day 3 (50% cell density). ‘Count’ is the number of genes enriched in a term. ‘Gene ratio’ is the percentage of total DEGs in the given term.

### Analysis of Alternative Splicing Events Linked to AGO2

Nuclear enrichment of AGO2 expands the possibilities for mechanisms of gene regulation to mechanisms that are distinct from the canonical translational silencing observed in the cytoplasm. Some of these nuclear-specific functions include regulation of transcription, chromatin organization, and alternative splicing [15]. As part of the pipeline developed on g.nome, we incorporated the tool rMATS to quantify the alternative splicing events. Custom code was written to facilitate visualization, including stacked bar plots and Venn diagrams comparing data from days 3, 5, and 7 as cell density increased.

We examined the five most common canonical alternative splicing events, including skipped exons, alternative 5’ splice sites, alternative 3’ splice sites, mutually exclusive exons, and retained introns (**Fig. 5A**). Relative to day 3, changes in all five categories of splicing events were observed in cells grown to greater confluence after 5 or 7 days (**Fig. 5B**). The majority of the altered splicing events are skipped exons and retained introns. We observed considerable overlap in events noted at day 5 and day 7 for the entire set of altered splicing events but there were also unique events observed at day 5 and 7 (**Fig. 5C**).

**Figure 5.**
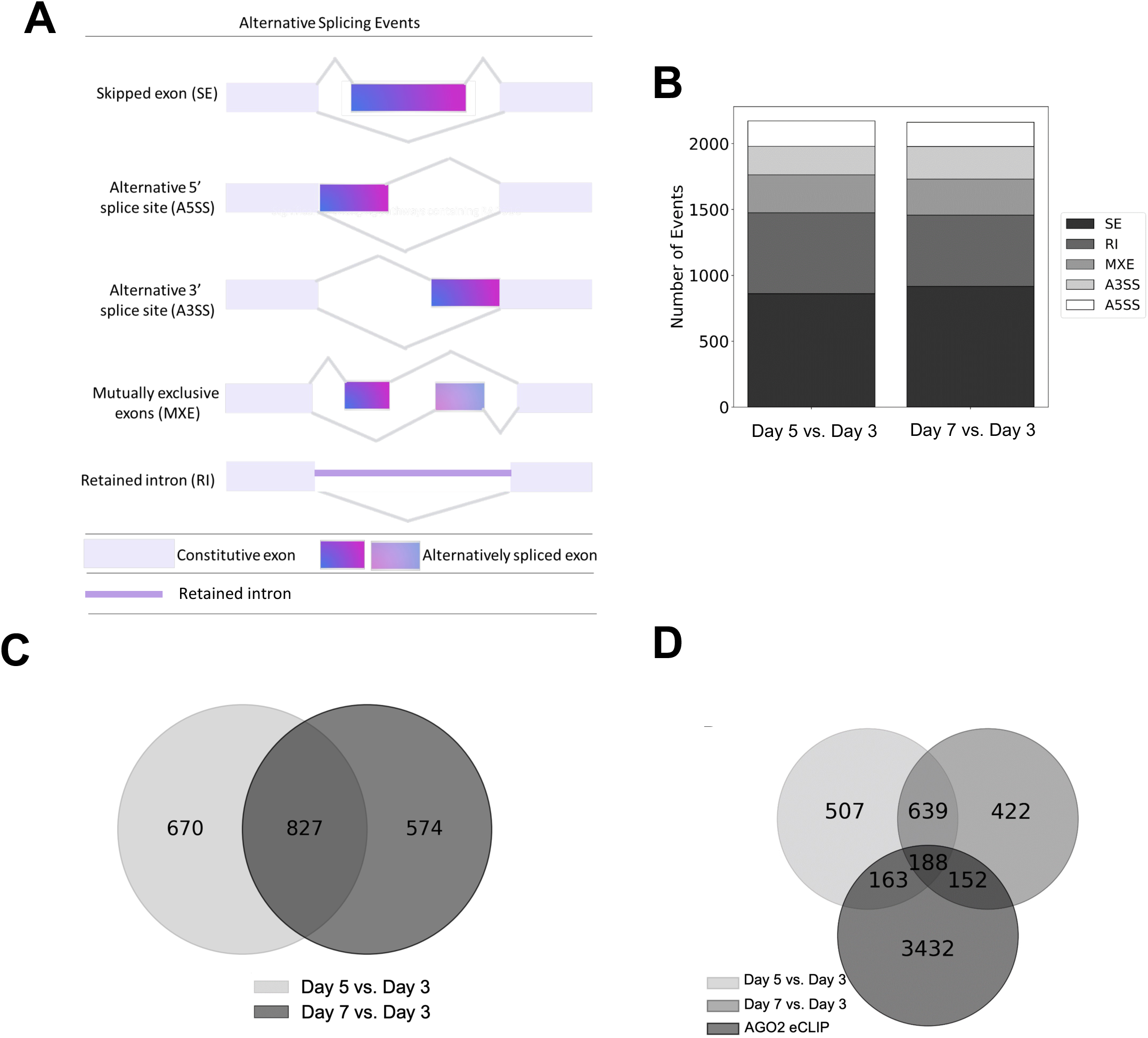
Differential alternative splicing events when AGO2 shuttles to nucleus. (A) Summary of the five different alternative splicing events detected with rMATS. (B) Bar graph showing number and type of splicing events detected on Day 5 (250% cell density) relative to Day 3 (50% cell density) and Day 7 (400% cell density) relative to Day 3 (50% cell density). (C) Venn diagram showing overlap of all alternative splicing events occurring on Day 5 and Day 7, both relative to Day 3. (D) Venn diagram showing overlap of genes undergoing alternative splicing events on Day 5 and Day 7 with genes associated with AGO2 from AGO2-eCLIP binding sites.

We also examined changes in splicing events at sites that previous studies have shown bind AGO2 [14]. We harnessed g.nome’s integrated Jupyter notebooks to combine outputs from our alternative splicing pipeline with additional AGO2 eCLIP data [18] enhancing our analysis of post-transcriptional gene regulation. By employing Python scripts within these notebooks, we were able to programmatically parse and annotate alternative splicing events, and subsequently align these with AGO2 eCLIP binding sites. This method facilitated an efficient and reproducible workflow, as the notebooks allowed for direct manipulation of data and visualization of results. This analysis showed that there was substantial overlap between genes at day 5 and day 7 that share altered splicing and have AGO2 bound near the relevant splice junctions (**Fig. 5D**). RNA sequences that bind AGO2 and have changes in splicing at Day 5 and Day 7 are candidates for modulation of splicing by AGO2:miRNA complexes.

### Proximity of AGO2 Binding Sites to Alternative Splicing Events

The shift of AGO2 localization from cytoplasm to nucleus can influence endogenous miRNA regulation [8]. This leads to the potential for dysregulation of miRNA targets both directly and indirectly, so we hypothesized that alternative splicing events that are most likely to be the result of direct AGO2 regulation should correlate with splice junctions that are near physically bound by AGO2 as determined by eCLIP [18].

We refined our analysis to alternative splicing events near AGO2 binding sites because these sites have the highest potential to affect splicing (**Fig. 6A**). We generated a histogram to represent the distribution of distances between identified alternative splice sites and AGO2 eCLIP sites (**Fig. 6B**). Splicing analysis was rMATS using a p-value cutoff of 0.05, with all splicing events analyzed. Analysis used both junction and exon counts. We identified genes with alternative splice sites in close proximity to AGO2 eCLIP sites, suggesting a potential mechanistic link between AGO2 binding and alternative splicing regulation.

**Figure 6.**
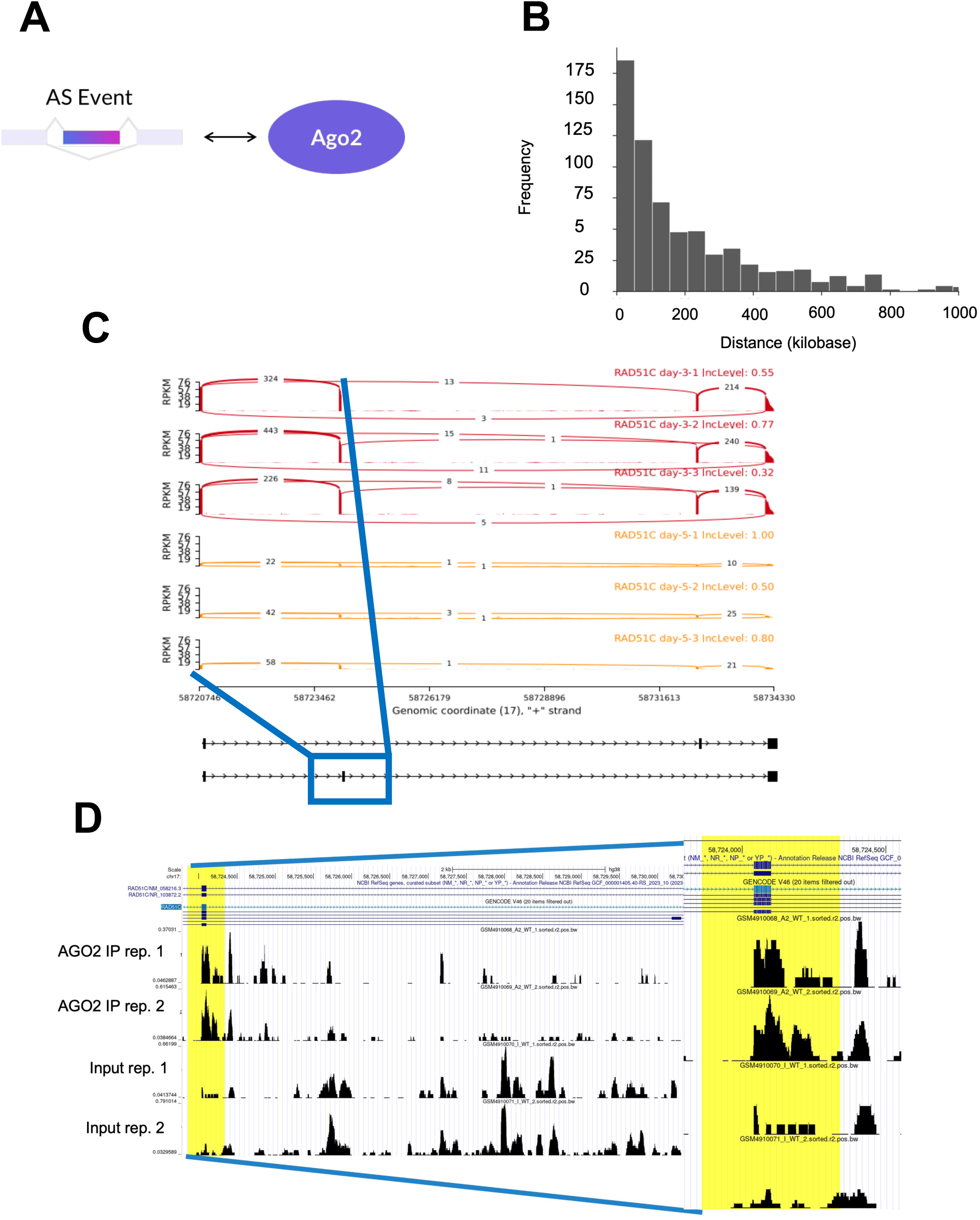
Top differential alternative splicing events refined based on proximity of splice junctions to AGO2-eCLIP bound sites. (A) Scheme showing selection of candidate alternative splicing events that were closest to AGO2 binding sites in the nucleus. (B) Histogram showing distribution of the distance between exon/intron junctions in genes undergoing alternative splicing and sites of AGO2 occupancy in the nucleus, as measured from AGO2-eCLIP-seq. (C) Sashimi plot of one representative candidate gene, RAD51, with skipped exon close to region of AGO2 occupancy. (D) AGO2 protein binding cluster within RAD51C exon-intron junction where the detected skipped exon event takes place.

We focused on RAD51 gene which exhibited an exon skipping event in proximity to an AGO2 binding site and observed a striking change in alternative splicing in day 5 relative to day 3 (**Fig. 6C**). These observations suggested that it is a good candidate for the control of splicing AGO2-mediated mechanisms.

### Comparison to other RNAseq-based differential expression and alternative splicing analysis tools

To validate the results of analysis with g.nome pipeline, we performed a comparative differential gene expression analysis using differential abundance nf-core pipeline [19] (**Sup. Fig. 3).** MDS analysis shows that replicates of datasets (Day 3, Day 5, Day 7) cluster together (**Sup. Fig. 3A**). Both pipelines agree on the global gene expression trends and show significant overlap between differentially expressed genes (**Sup. Fig. 3B**). We observed larger portion of genes showing upregulation rather than downregulation in Day 5 (250% confluence) or Day 7 (400% confluent) as compared to Day 3 (50% confluence) (**Sup. Fig. 3B**).

## Discussion

### The need for transparent pipelines for the analysis of large datasets

Modern technologies for nucleic acid sequencing generate vast amounts of data. This data enables insights into basic biological pathways and disease. The challenge, however, is how to sift through large datasets to fully exploit their content and avoid being drawn to conclusions that are not rigorously supported.

Reproducibility and accessibility form the cornerstone of scientific discovery, and these principles apply to the interpretation of large datasets. Unlike older forms of data that could be viewed on a micrograph or gel and grasped immediately, datasets require careful analysis. This analysis can be difficult for scientists who are primarily experimentalists to use them properly and fully exploit their potential. For example, experiments are time-consuming, and it may not be straightforward for a non-computational biologist to sift through “statistically significant” data and identify the leads that are most worthy of in-depth investigation. It is important to reduce the “black box” nature of computational analysis as much as possible to that users can better understand their data, use it to design experiments, and provide transparent explanations for how the data was used.

### g.nome, a platform to facilitate data analysis for experimental biologists

The goal of g.nome is accessible, thorough, and integrated data analysis, bridging the gap between complex datasets and meaningful biological insights through g.nome’s Canvas and Guided Workflows. Here we describe a pipeline integrating gene expression and splicing data. Our analysis illustrates g.nome’s capability to synthesize complex analyses for researchers without computational backgrounds into a visual node-and-wire language. This integration is not trivial; it encompasses multiple layers of biological regulation and interaction that are pivotal for understanding cellular responses and disease mechanisms. By allowing researchers to navigate through these layers without the need for extensive computational training, g.nome democratizes the exploration and interpretation of biological data.

As analysis techniques become more advanced, incorporating elements from artificial intelligence and systems biology to tackle predictive modeling and the dynamic interactions within biological networks, g.nome is poised to adapt and incorporate these methodologies. Its design philosophy centers on democratizing access to sophisticated analyses, allowing researchers to focus on scientific inquiries rather than computational complexities.

g.nome’s trajectory mirrors the broader evolution in biological data analysis by emphasizing usability, integration, and adaptability. It aims to empower researchers to harness the full potential of their data, enabling a deeper exploration of gene regulation, cellular processes, and, ultimately, the mechanisms underlying health and disease. As the field evolves, g.nome’s ability to incorporate new analytical paradigms and data types will make it an essential tool in the biologist’s arsenal, facilitating discoveries that were once beyond reach.

### g.nome features

g.nome offers a cloud-based graphical platform enabling scientists to conduct sophisticated analyses without extensive computational expertise or setting up complex computation environments. This platform integrates a comprehensive toolkit for autonomous analysis, featuring a user-friendly graphical interface for straightforward job execution and intuitive “drag and drop” dataset analysis.

The platform addresses the challenges faced by users with limited bioinformatics knowledge. Tasks such as aligning sequencing data to the human genome, identifying various transcripts, and comparing datasets can be daunting. The inclusion of a splicing module simplifies these processes by quickly generating analyses, quality control metrics, and summary graphs directly from the platform’s splicing package, eliminating the need to navigate through Excel spreadsheets.

To this end, we have introduced an alternative splicing workflow developed on g.nome, a cloud-based graphical platform that empowers experimental biologists to perform complex analyses without necessitating a deep understanding of computational tool setup. g.nome integrates an array of contemporary tools necessary for independent analysis, providing a user-friendly graphical interface that simplifies the running of jobs and allows for intuitive “drag and drop” analysis of various datasets.

The platform helps solve the problem faced by biologist with little bioinformatic expertise. Running analysis programs to align sequencing data to the human genome, characterizing different transcripts in the genome, aligning different sample datasets, can be challenging. It is convenient to have a splicing module that could quickly generate analysis, quality control metrics, and summary graphs. Data is provided automatically from the platform splicing package. The package also does differential analysis for sample versus control group, which avoids requiring the user to run another program that requires some coding expertise.

### Application of g.nome to the impact of nuclear AGO protein

The goal of g.nome is to enable non-computational biologists to independently ask questions of their data. Our research has harnessed the capabilities of g.nome to uncover the dynamics of gene expression and alternative splicing in response to changes in cellular density and AGO2 localization. We observed that as cells of the colorectal cancer line HCT116 grow to high density, AGO2, a key player in RNA-induced silencing complex, shifts to the nucleus [8]. This redistribution is hypothesized to affect the recognition of intronic sequences by miRISC, potentially influencing alternative splicing events.

The g.nome platform has enabled the straightforward identification of splicing changes. With g.nome, the complexities of bioinformatics are distilled into a transparent and straightforward process, enabling rapid progression from dataset comparison to experimental tests. This not only underscores the regulatory role of AGO2 in alternative splicing but also highlights the potential of g.nome to serve as a valuable tool in the broader scientific community.

The exploratory analysis using custom code goes even further to incorporate a large dataset from eCLIP to identify AGO2 binding sites that are close to sites of significant splicing change. This was done systematically across all alternative splicing events. It also produced Sashimi plots that make the data more transparent and interpretable. Using the g.nome custom code we learned that there was a high frequency of alternative splicing events that were close in distance to AGO2 occupancy sites. This analysis provides us with candidate lists to validate experimentally.

## Conclusion

The landscape of biological data analysis is undergoing a profound transformation, driven by the exponential growth of data and the increasing complexity of biological questions. As the volume and variety of data expand, from single-cell genomics to complex proteomic patterns, researchers are seeking more sophisticated tools to navigate and interpret this wealth of information. This evolution is marked by a shift towards integrative approaches that can handle the intricacy and interconnectedness of biological systems.

In this changing landscape, platforms like g.nome are set to play a pivotal role. By offering a suite of tools designed for the modern experimental biologist, g.nome aligns with the trend towards more accessible, comprehensive, and integrated data analysis. It stands at the intersection of innovation and practicality, providing a bridge between complex data and actionable biological insights. As an example, we describe a pipelines that combines data on gene expression and splicing. These datasets are complex and their combination poses computation and analytical challenges. g.nome enabled construction of a straightforward pipeline that enabled efficient independent analysis by non-computational biologists.

## Data availability

The small-RNA and whole transcriptome RNA-sequencing data has been deposited in NCBI’s Gene Expression Omnibus and is accessible through GEO Series accession number GSE236946. AGO2-eCLIP sequencing data was previously published (14).

## Author contributions

D.R.C. wrote the manuscript and supervised the experiments. K.C.J. performed experiments. K.C.J., R.B. analyzed data using g.nome. M.K. developed the g.nome pipeline. X.K. performed RNA-seq analyses using nf-core pipeline. K.C.J., R.B., M.K. assisted in writing the manuscript.

## Supplementary Information

### RNAseq-based differential expression and alternative splicing analysis using g.nome - code

*Preprocessing and Quality Control*

fastp --detect_adapter_for_pe --trim_poly_g --in1 WCDay7Rep2_S60_R1_001.fastq.gz --in2 WCDay7Rep2_S60_R2_001.fastq.gz … --thread 2 --out1 WCDay7Rep2_S60_R1_001.fastq.gz --out2 WCDay7Rep2_S60_R2_001.fastq.gz

fastqc WCDay3Rep1_S53_R1_001.fastq.gz WCDay3Rep1_S53_R2_001.fastq.gz WCDay3Rep2_S54_R2_001.fastq.gz … --threads 3 --outdir output-dir

multiqc --outdir output input_dir

*Read Alignment*

STAR --readFilesIn WCDay7Rep2_S60_R1_001.fastq.gz

WCDay7Rep2_S60_R2_001.fastq.gz --genomeDir reference_dir --readFilesCommand zcat --outSAMtype BAM Unsorted --runThreadN 10 --outFileNamePrefix input_dir

*BAM File Processing*

picard FixMateInformation -ADD_MATE_CIGAR -INPUT Aligned.out.bam

picard MarkDuplicates --INPUT Aligned.out.bam --METRICS_FILE metrics.txt -- OUTPUT Aligned.out_marked.bam --TMP_DIR tmp

*RNA-seq Metrics Calculation*

rnaseqc GRCh38_gencode.v45.annotation.gtf.gz_collapsed.gtf

Aligned.out_marked.bam input_dir

*Differential Expression Analysis*

Rscript DEseq2.R --show-rownames --show-colnames --gene-count-matrix counts_deseq2.txt --sample-metadata sample_metadata.txt --control-name d3 --p-value 0.05 --fc-cut-off 0.75 --output-dir output-dir

*Gene Set Enrichment Analysis (GSEA)*

Rscript GSEA.R --control-name d3 --dds dds_object.rds --fc-cut-off 0.75 --p-value 0.05 - -organism human --output-dir output-dir

**Supplementary Figure 1.**
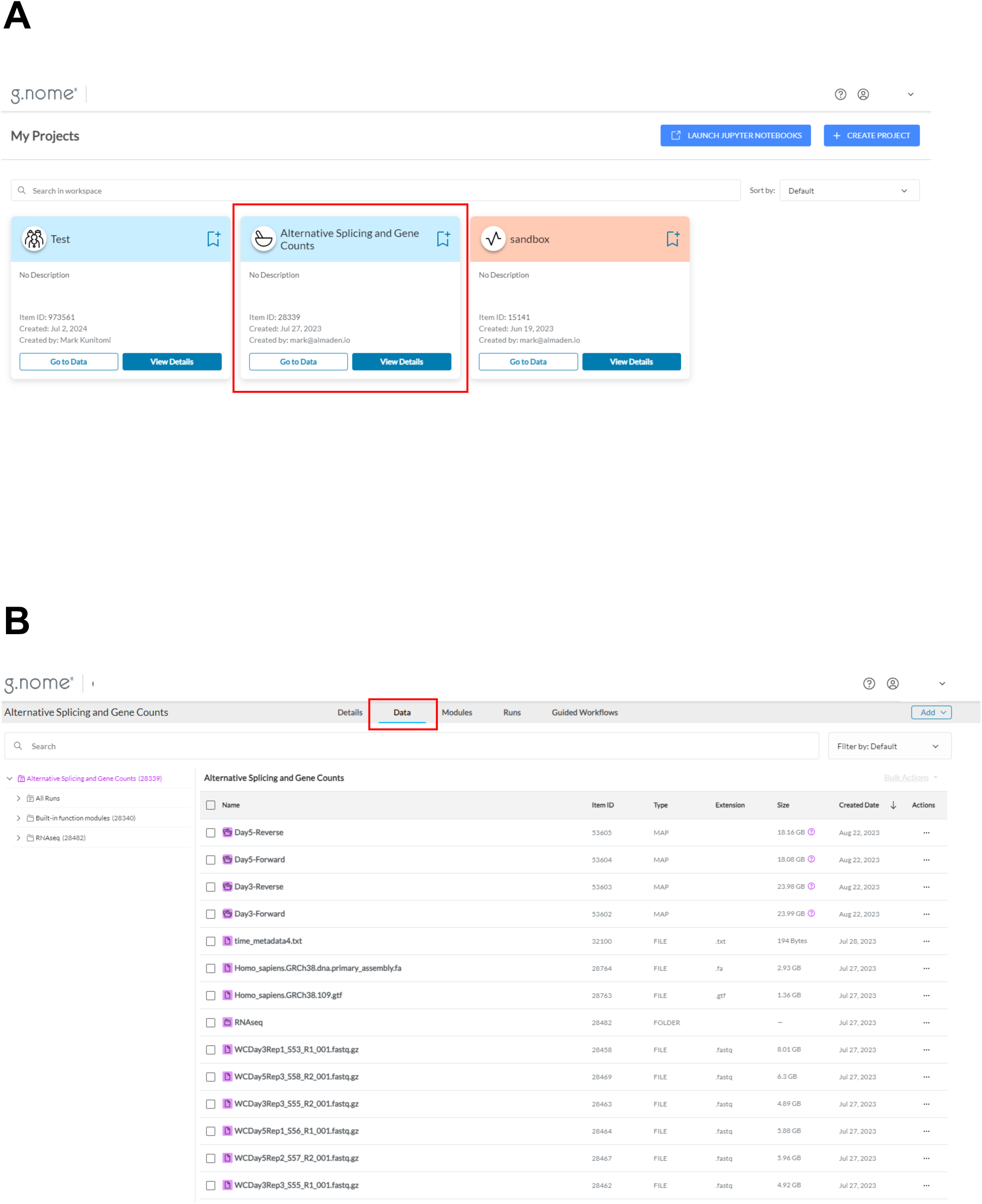
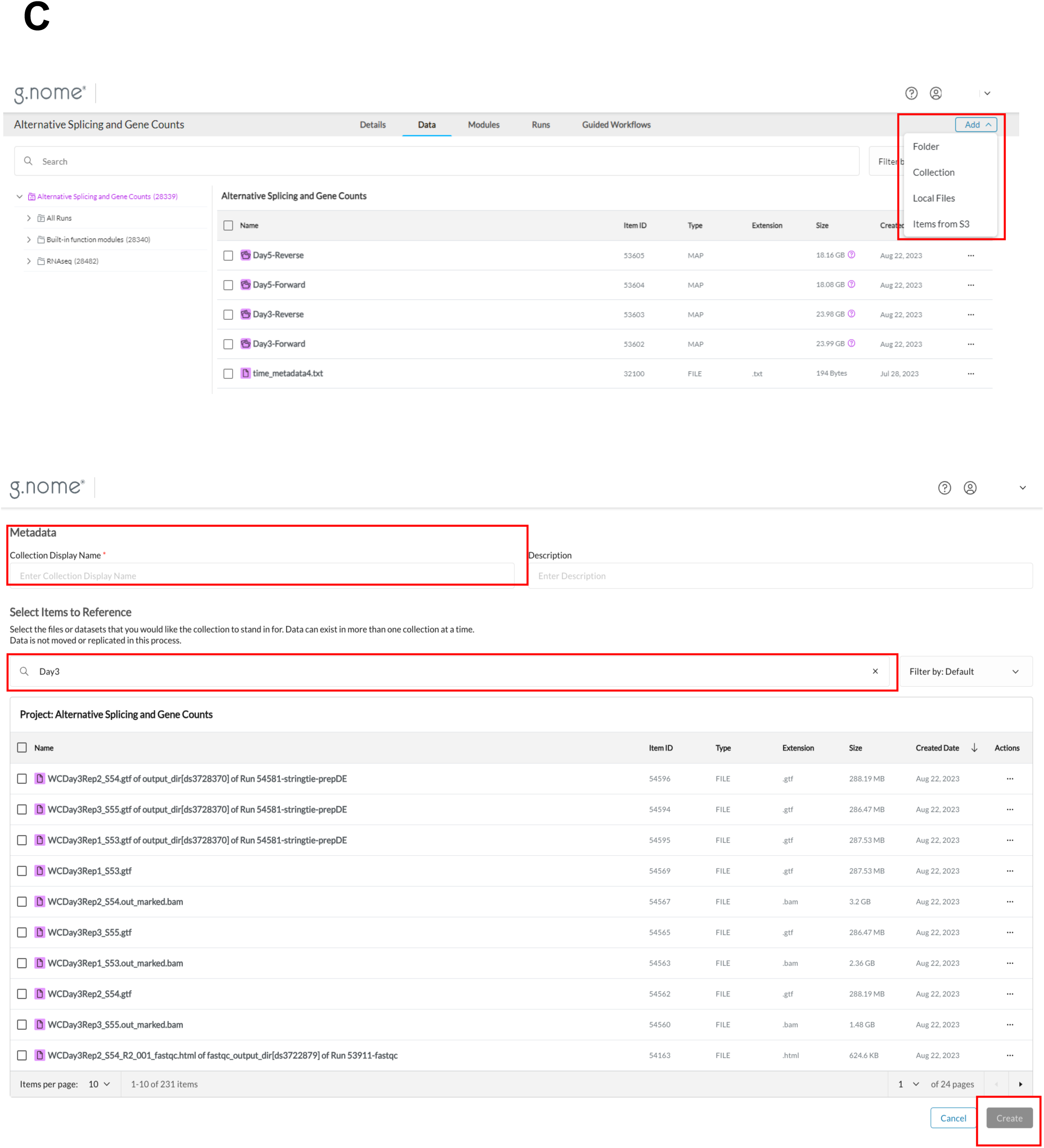
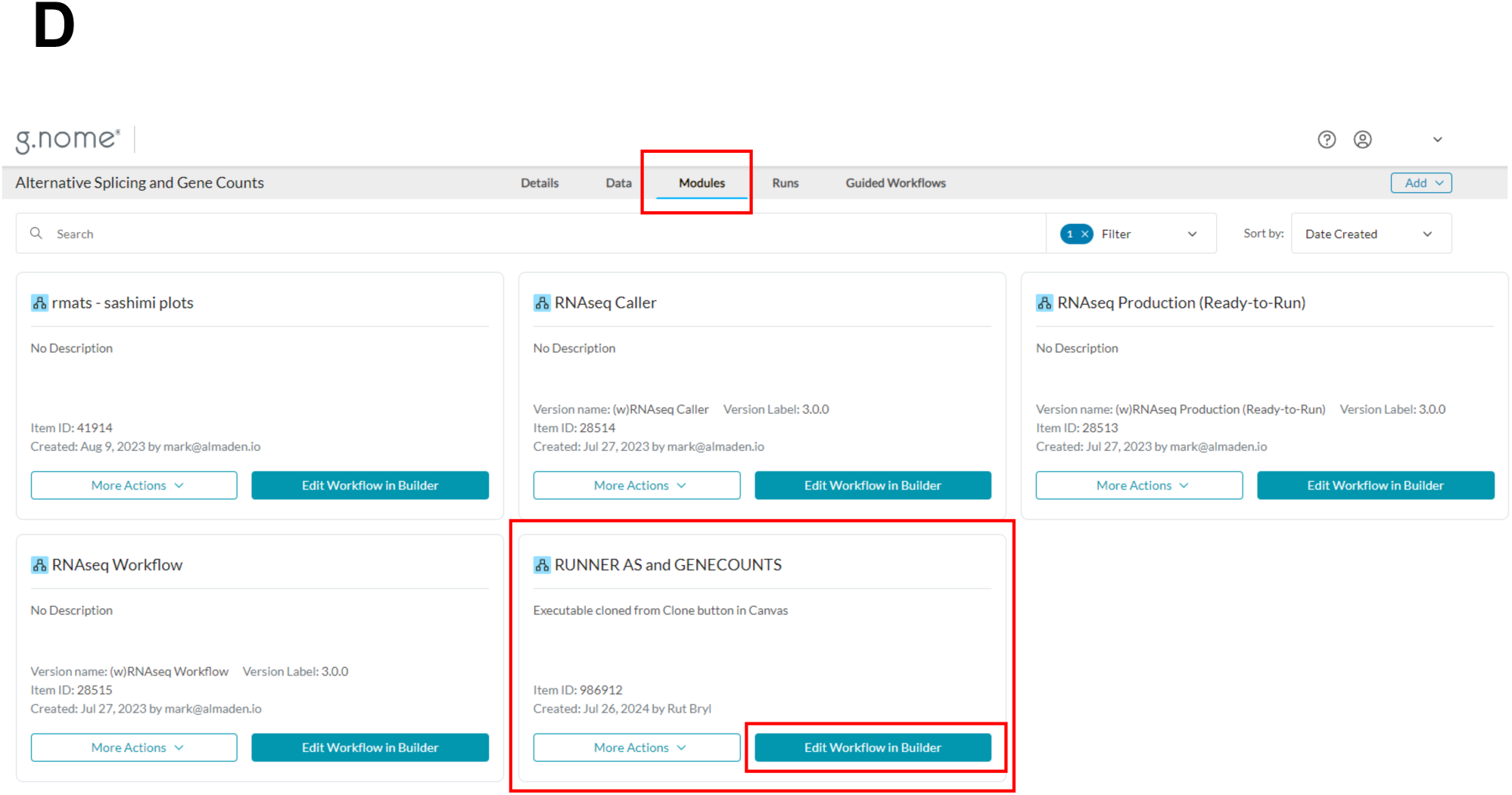
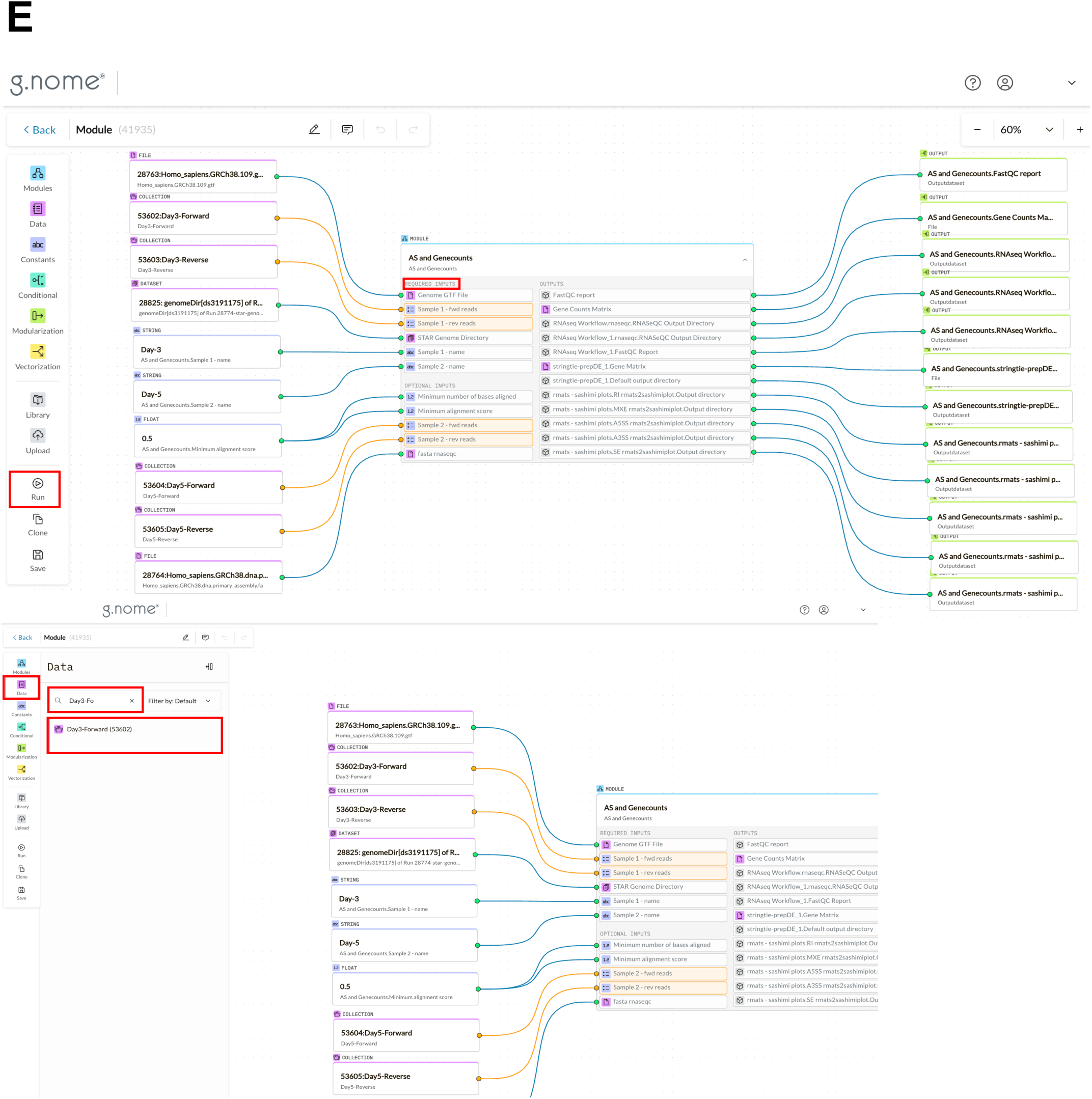
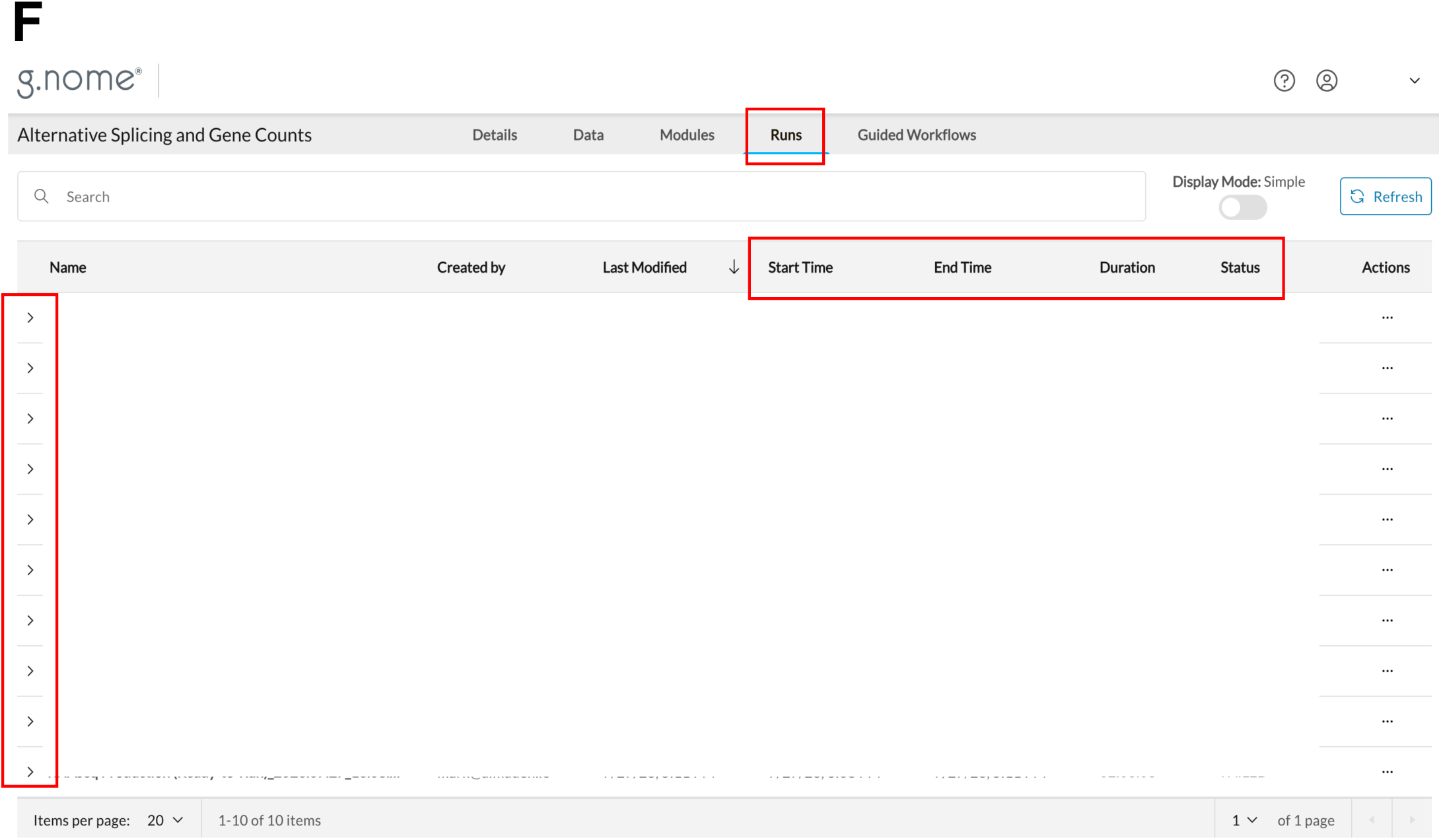
Usage of g.nome pipeline for alternative splicing analysis by non-bioinformaticians. (A) User can create and choose projects according to their needs. For this analysis, “Alternative Splicing and Gene Counts” was selected. (B) The raw sequencing files and human genome assembly can be uploaded into the “Data” folder under a given project (in this case “Alternative Splicing and Gene Counts”. (C) User can create a batch collection of relevant data using “Add” and “Collection” function. The search bar can be used to look up the relevant files. (In this example, “Day3-Forward” collection was created, and the search bar was used to look up the relevant files with “Day3” and “R1” in the file names for forward reads for each of the three biological replicates from the Day 3 time point.) (D) After all batch collections were created, a desired module is selected to perform the analysis. In this example, the alternative splicing pipeline module, “Runner AS and Genecount,” was chosen and “Edit Workflow in Builder” was selected. (E) Within the splicing module, there are empty cells for “requested inputs”. The circle on the left side of “Sample 1 – name” is dragged to the left into the open workspace area. The “String Value” can be edited to be the sample name (in this example “Day3” and “Day 7”). Then, on the right side of the screen, “Data” is selected, and the search bar can be used to look up the batch collections. User can click and drag and drop the batch collections into the right-hand workspace (in this example “Day3-Forward” batch collection). To designate the input for sample 1 on the splicing module, the orange circle on the “Sample 1 – fwd reads” required input is dragged to connect to the open white circle on forward reads data collection (in this example “Day3-Forward”). This is repeated for the reverse collection and for forward and reverse collections for “Sample 2” inputs. Finally, on the right side panel, the “Run” button is selected. (F) The edited workflow can be saved, and the progress of the run can be monitored within the “Runs” tab at the top center panel. The down arrow on the run can be selected to reveal a dropdown menu with output files such as quality control measurements and summary graphs displaying formatted data.

**Supplementary Figure 2.**
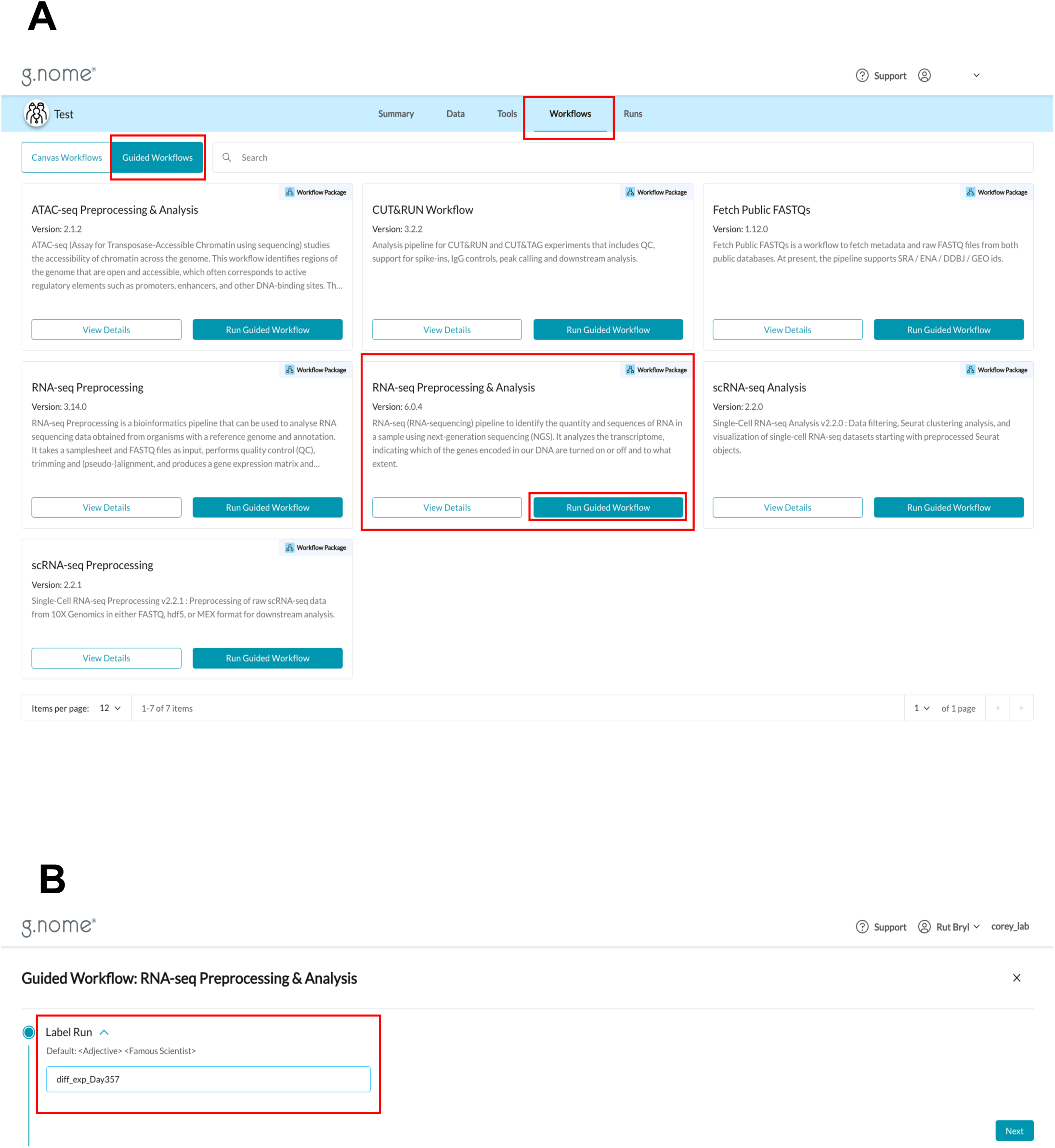
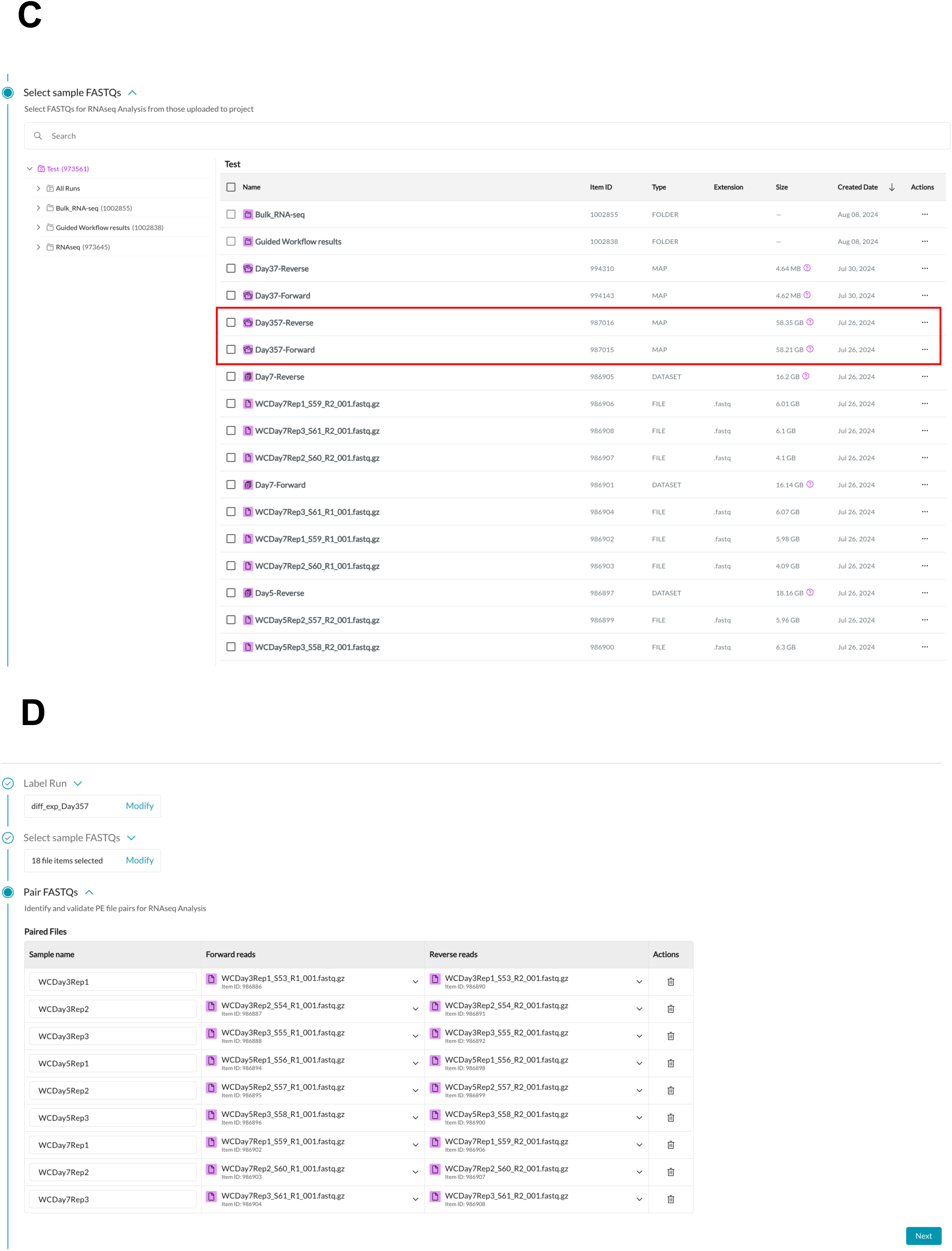
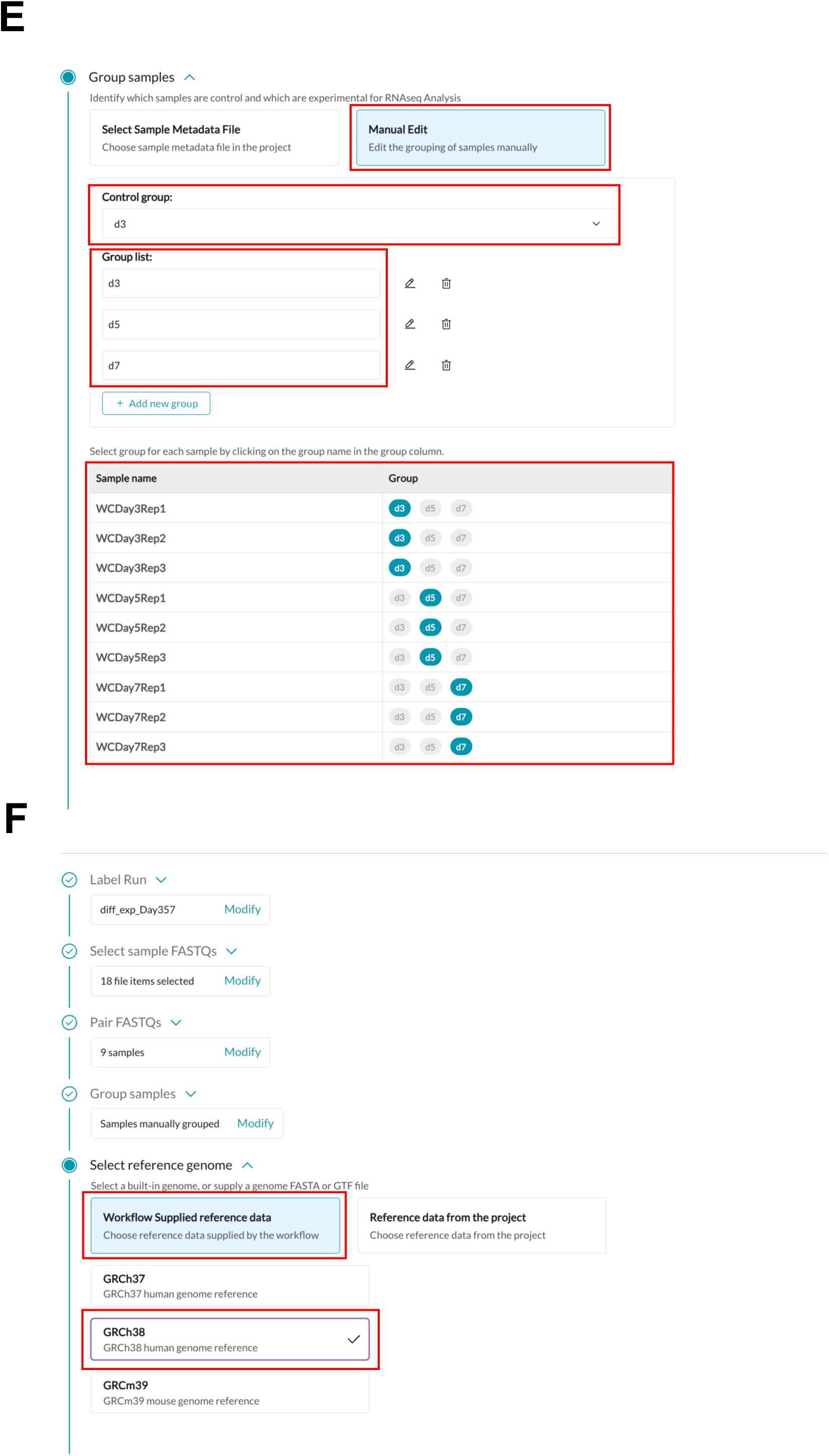
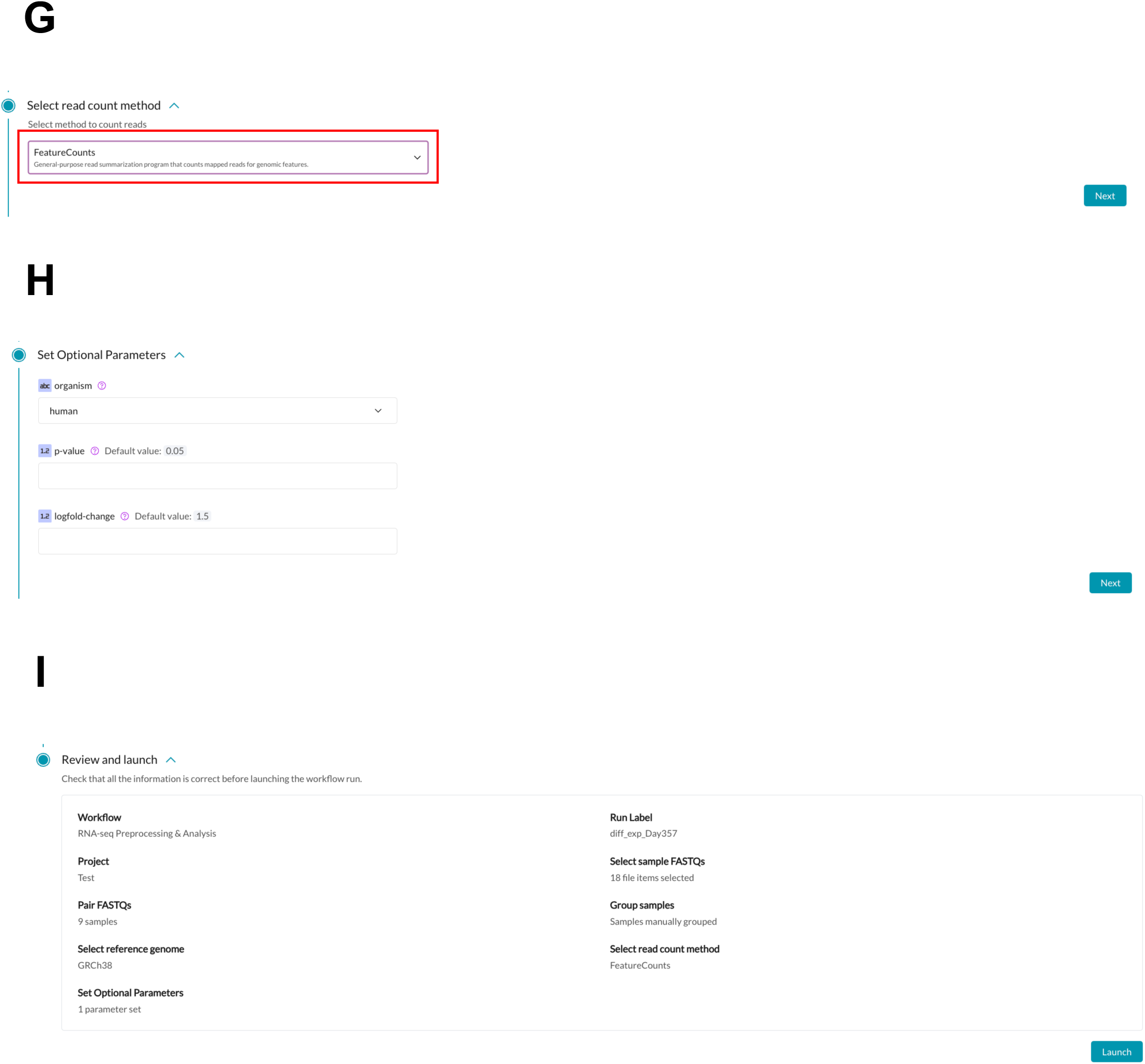
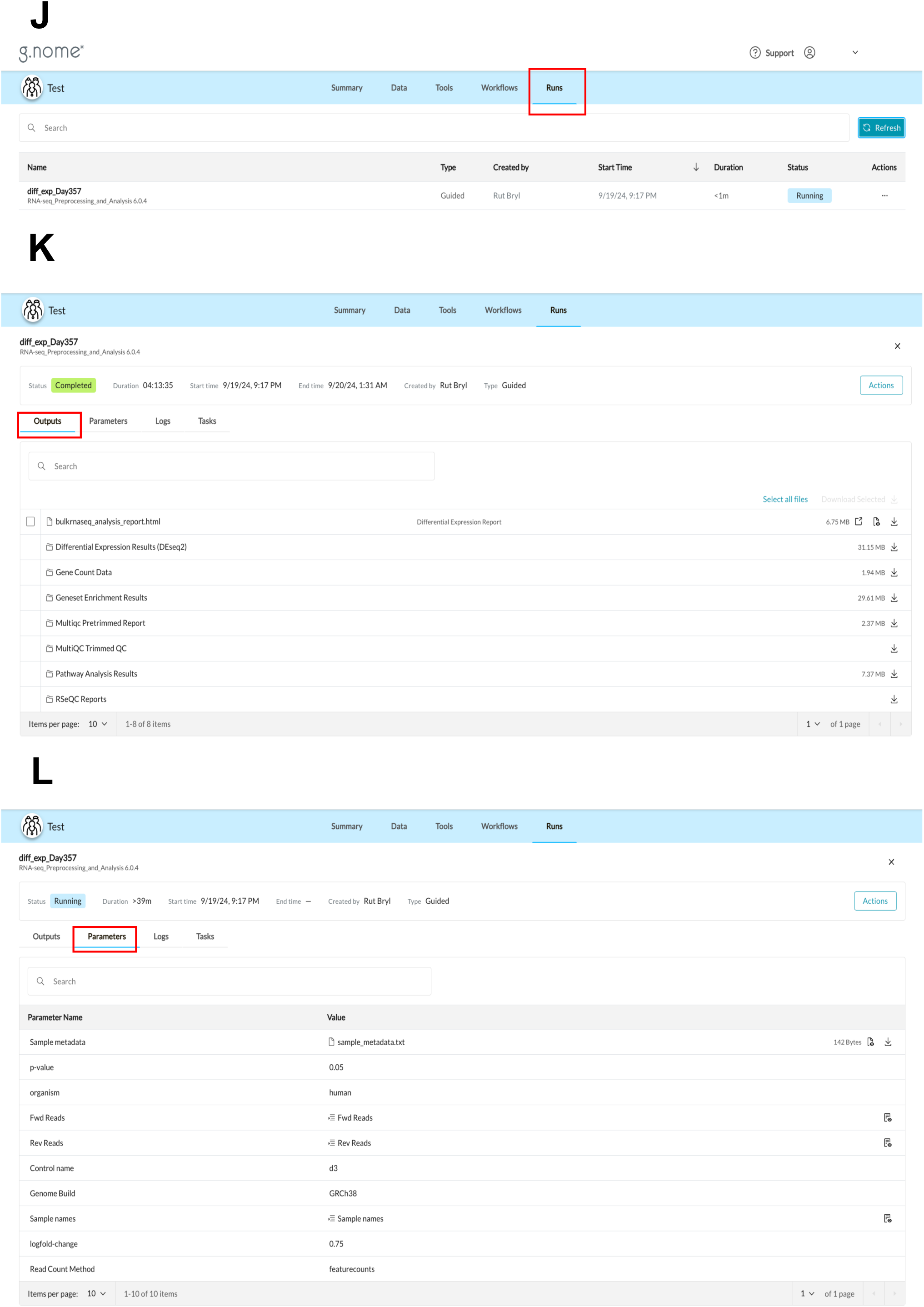
Usage of g.nome pipeline for differential gene expression analysis by non-bioinformaticians. (A) Under “Workflows” tab, “Guided Workflows” button was clicked and the differential gene expression pipeline module “RNA-seq Preprocessing & Analysis” and “Run Guided Workflow” was selected. (B) The run was labeled and (C) collections “Day357-Forward” and “Day357-Reverse” were selected as sample FASTQs. (D) The FASTQ files were then paired, according to their biological replicates. (E) Next, the samples were grouped – “Manual Edit” button was selected, and control group and group lists were defined. Finally, samples were matched with their respective group. (F) Reference genome was selected as “Workflow Supplied reference data” and “GRCh38”. (G) A read count method was “FeatureCounts”. (H) Optional parameters were left as default. (I) Before launching the run, the user is provided with the summary of the chosen parameters. (J) The run can be monitored in the “Runs” tab. (K) The name of the run was then clicked, which takes the user to the outputs, where analyzed data can be accessed. (L) Basic parameters such as p-value, log fold-change, reference genome as well as sample metadata and FASTQ files used can be viewed in “Parameters” tab.

**Supplementary Figure 3.**
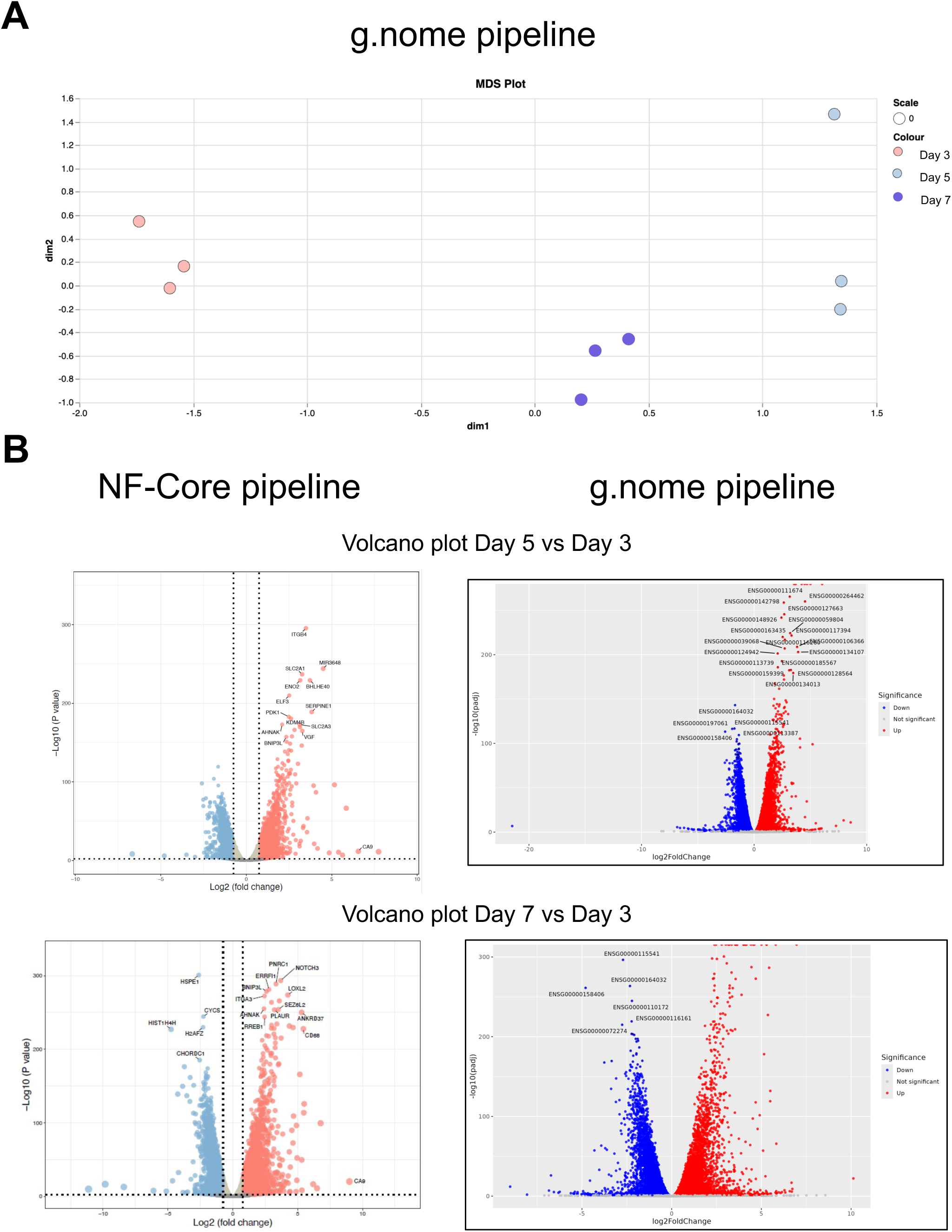
Comparison of differential gene expression analysis using g.nome or nf-core. (A) Multidimensional-scaling (MDS) analysis of three replicates per cell density condition shown: Day 3 (50% cell density), Day 5 (250% cell density), and Day 7 (400% cell density) prepared using g.nome. (B) Volcano plots showing genes significantly downregulated and upregulated in Day 5 (250% cell density) relative to Day 3 (50% cell density) and Day 7 (400% cell density) relative to Day 3 (50% cell density).

